# Effects of non-invasive brain stimulation on effective connectivity during working memory task in Neurofibromatosis Type 1 patients

**DOI:** 10.1101/2024.10.16.618671

**Authors:** Marta Czime Litwińczuk, Shruti Garg, Stephen R. Williams, Jonathan Green, Caroline Lea-Carnall, Nelson J Trujillo-Barreto

## Abstract

This study examined the effects of anodal transcranial direct current stimulation (atDCS) on effective connectivity during a working memory task. Eighteen adolescents with Neurofibromatosis Type 1 (NF1) completed a single□blind sham□controlled cross□over randomised atDCS trial. Dynamic causal modelling was used to estimate the effective connectivity between regions that showed working memory effects from the fMRI. Group-level inferences for between sessions (pre- and post-stimulation) and stimulation type (atDCS and sham) effects were carried out using the parametric empirical Bayes approach. A correlation analysis was performed to relate the estimated effective connectivity parameters of left dlPFC pre-atDCS and post-atDCS to the concentration of gamma-aminobutyric acid (GABA) measured via magnetic resonance spectroscopy (MRS-GABA). Next, correlation analysis was repeated using all working memory performance and all pre-atDCS and post-atDCS connectivity parameters. It was found that atDCS decreased average excitatory connectivity from left dorsolateral prefrontal cortex (dlPFC) to left superior frontal gyrus and increased average excitatory connectivity to left globus pallidus. Further, reduced average intrinsic (inhibitory) connectivity of left dlPFC was associated with lower MRS-GABA. However, none of the connectivity parameters of dlPFC showed any association with performance on a working memory task. These findings suggest that atDCS reorganised connectivity from frontal to fronto-striatal connectivity. As atDCS-related changes were not specific to the effect of working memory, they may have impacted general cognitive control processes. In addition, by reducing MRS-GABA, atDCS might make dlPFC more sensitive and responsive to external stimulation, such as performance of cognitive tasks.

**Highlights:**

- atDCS was applied to left dlPFC in NF1 patients during working memory
- After atDCS, no effect on modulatory connectivity
- Evidence for increased N-back average connectivity from dlPFC to globus pallidus
- Less dlPFC MRS-GABA was associated with less dlPFC inhibition

## Introduction

Neurofibromatosis 1 (NF1) is a single-gene autosomal dominant neurodevelopmental disorder with birth incidence of 1:2700 Evans, Howard [1]. The condition is caused by mutation of the NF1 gene, encodes the neurofibromin protein, and regulates the Ras-MAPK molecular pathway [2, 3]. Physiologically, NF1 is associated with skeletal abnormalities, brain and peripheral nerve tumours [4]. In addition, NF1 patients tend to underperform in academic settings compared to their siblings without an NF1 diagnosis, including poorer performance on assessments of intelligence, visuospatial skills, social competence, executive function and attention [5]. Notably, Shilyansky, Karlsgodt [6] found that NF1 patients score lower on working memory tasks than controls and that they are more sensitive to the effects of increased memory load than controls. In addition, Shilyansky, Karlsgodt [6] explored the neural substrates of these deficits using functional magnetic resonance imaging (fMRI) and demonstrated that deficits in working memory of NF1 patients were associated with hypoactivity in dorsolateral prefrontal cortex (dlPFC), frontal eye fields, and parietal cortex, as well as striatal regions. Thus, the working memory deficits in NF1 patients may be explained by disrupted activity of left frontal regions associated with working memory.

To address these physiological and psychosocial alterations, clinical neuroscience has explored therapeutic effects of non-invasive brain stimulation (NIBS). Application of NIBS has shown some success in improving cognitive functioning in children and adolescents with neurodevelopmental conditions, such as autism spectrum condition and attention deficit hyperactivity disorder [7-9]. Garg, Williams [10] investigated effects of NIBS on working memory performance in NF1 patients. Anodal transcranial direct current stimulation (atDCS) was applied to left dlPFC during performance of a working memory task [11, 12]. atDCS is thought to increase cortical excitability through depolarisation of the resting membrane potential. Garg, Williams [10] demonstrated that application of atDCS was associated with decreased levels of GABA in the dlPFC, as measured with magnetic resonance spectroscopy (MRS). However, atDCS had transient effects on blood oxygen level dependent (BOLD) signal in fMRI data, and it did not influence behavioural outcomes.

To further understand the effects of atDCS on NF1 patients it is important to understand how brain regions coordinate their activity with each other during performance of working memory tasks prior and post NIBS. Here, we further analyse the data obtained by Garg, Williams [10], as we investigate the effects of a sham-controlled anodal atDCS administration trial on effective connectivity of left dlPFC in adolescent patients with NF1. To this end, this work will employ dynamic causal modelling (DCM) [13]. Since in the original Garg, Williams [10] study the administration of atDCS aimed to increase dlPFC excitability, we hypothesise that the administration of atDCS will result with a reduction of inhibitory intrinsic connectivity (self-connectivity) of left dlPFC, which will increase connectivity from left dlPFC to other regions involved with working memory. We further hypothesise that effective connectivity parameters of left dlPFC will show a positive correlation with GABA concentration and working memory performance.

## Methods

### NF1 Participants

The data used in this study has been previously described in Garg, Williams [10]. Thirty-one adolescents (16 males, 15 females) aged 11–17 years were recruited via the Northern UK NF-National Institute of Health diagnostic criteria [National Institutes of Health Consensus Development Conference. Neurofibromatosis conference statement. Arch. Neurol. 45, 575–578 (1988).] and/or molecular diagnosis of NF1; (ii) No history of intracranial pathology other than asymptomatic optic pathway or other asymptomatic and untreated NF1-associated white matter lesion or glioma; (iii) No history of epilepsy or any major mental illness; (iv) No MRI contraindications. Participants on pre-existing medications such as stimulants, melatonin or selective serotonin re-uptake inhibitors were not excluded from participation. The study was conducted in accordance with local ethics committee approval (Ethics reference: 18/NW/0762, ClinicalTrials.gov Identifier: NCT0499142.

Registered 5th August 2021; retrospectively registered, https://clinicaltrials.gov/ct2/show/NCT04991428). All methods were carried out in accordance with relevant guidelines and regulations.

### Study design

The impact of atDCS on working memory was assessed through a two-arm, single-blind (participant), sham-controlled cross-over study (Figure 1). Each participant attended two visits, spaced at least one week apart. During each visit they received either the atDCS or sham stimulation, with the order randomized and counterbalanced. Participants maintained their regular medication schedules, including stimulants. During the visits, participants were comfortably positioned in a scanner, and high-resolution T1-weighted images were acquired. Participants performed the N-back working memory task (Supplementary Material 1) during four 6-minute-long sessions (24 minutes in total) while fMRI data were collected.

**Figure 1.**
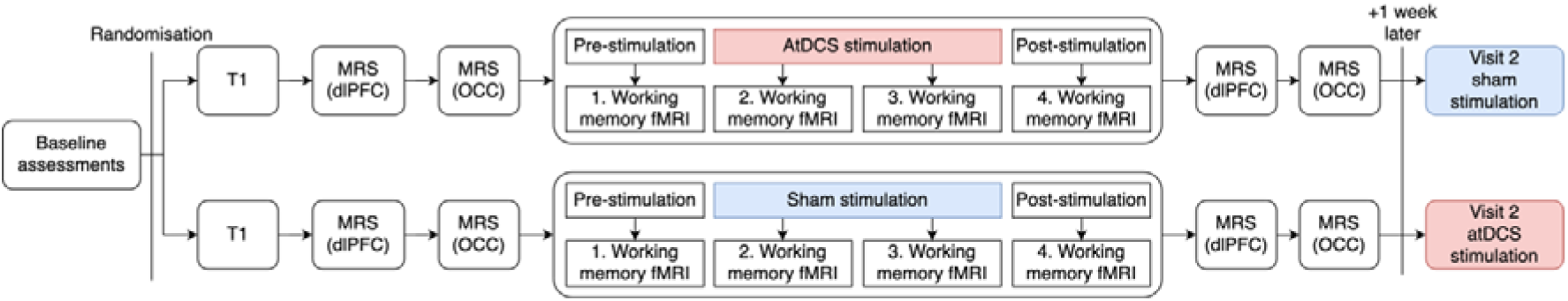
A schematic illustration of the study design and image acquisition. Depending on randomisation, during Visit 1, either atDCS or sham stimulation was applied during session 2 and 3 of working memory fMRI acquisition.

Stimulation (atDCS or sham) was applied for 15 minutes during sessions 2 and 3. Between each session, the participants were asked if they were comfortable, and instructions were repeated. Additionally, T2-weighted images were acquired during the first visit and evaluated by a paediatric neuroradiologist to exclude NF1-associated tumours.

### Stimulation

Stimulation was administered using a NeuroConn DC-STIMULATOR MR, with the anode positioned at F3 and the cathode at Cz according to the international 10–20 system. The scalp was first cleaned with Nuprepgel, and Ten20-paste was applied as a conductive medium between the scalp and the electrodes. During atDCS stimulation, the current was gradually increased over 15 seconds, maintained at 1 mA for 15 minutes, and then decreased over 15 seconds. During sham stimulation, the current was ramped up over 15 seconds and then immediately turned off. The parameters for the current were determined based on prior safety trials conducted with this cohort (clinical trials identifier: NCT03310996).

Stimulation was delivered via a NeuroConn DC-STIMULATOR MR with the anode placed over F3 position in the international 10–20 system and the cathode over the Cz position. Scalp was cleaned with Nuprepgel and Ten20-paste was used as a conductive medium between the scalp and the electrodes. For anodal stimulation, the current was ramped up over 15 s, held at 1 mA for 15 min and then ramped down over 15 s. For sham stimulation, the current was ramped up over 15 s and then immediately turned off. The current parameters were chosen based on our previous experience from a pilot clinical trial of safety in this cohort (clinical trials identifier: NCT03310996).

### Structural MRI and MRS acquisition

Scanning was conducted using a Philips Achieva 3 T MRI scanner (Best, NL) equipped with a 32-channel head coil. First, 3D T1-weighted magnetic resonance images were obtained in the sagittal plane with a magnetization-prepared rapid acquisition gradient-echo sequence (repetition time = 8.4 ms; echo time = 3.77 ms; flip angle = 8°; inversion time = 1150 ms; in-plane resolution = 0.94 mm; 150 slices with 1 mm thickness). Next, a T2-weighted structural scan was performed using a turbo spin echo sequence (TR = 3756 ms; TE = 89 ms; 40 slices of 3 mm thickness and 1 mm gap; in-plane resolution = 0.45 mm). Single-voxel ^1^H MRS data were collected before and after stimulation from two volumes of interest (VOI) in each participant: one VOI (40 × 20 × 24 mm) in the left dlPFC and a control VOI (20 × 50 × 20 mm) in the posterior occipital lobe (OCC), centred on the mid-sagittal plane to encompass both hemispheres. Water-unsuppressed spectra were recorded from these locations to serve as references. To detect GABA+ (so called, because the edited signal contains contributions from co-edited macromolecules as well as GABA), water-suppressed GABA-edited MEGA-PRESS spectra [14, 15] were acquired (repetition time = 2000 ms; echo time = 68 ms; 1024 sample points at a spectral width of 2 kHz, as detailed in Sanaei Nezhad, Anton [16]. The acquisition for the dlPFC MRS took approximately 7 minutes, with 96 averages, while the occipital voxel MRS took about 3 minutes, with 32 averages. The number of averages was adjusted to ensure comparable spectral quality between the DLPFC and OCC. Processing of MRS data is described in Supplementary Material 2.

### Functional MRI acquisition

Imaging was performed on a 3 Tesla Philips Achieva scanner using a 32-channel head coil with a SENSE factor 2.5. To maximise signal-to-noise (SNR), we utilised a dual-echo fMRI protocol developed by Halai, Welbourne [17]. The fMRI sequence included 36 slices, 64×64 matrix, field of view (FOV) 224×126×224 mm, in-plane resolution 2.5×2.5 mm, slice thickness 3.5 mm, TR=2.5 s, TE = 12 ms and 35 ms. The total number of volumes collected for each fMRI session was 144. Processing of fMRI data is described in Supplementary Material 3.

### Volumes of interest

To select volumes of interest (VOIs), we considered regions that have been previously involved with N-back task performance in adults [18] and adolescents [19]. In addition, during selection of VOIs we considered regions affected by working memory in this dataset. To identify these regions, first-level mass univariate analysis using the General Linear Model based on the canonical haemodynamic response function. A high-pass filter with a cut-off at 128 seconds was applied to remove slow signal drifts. An autoregressive model of order 1 was fitted to estimate and remove serial correlations [20]. The outlier volume censoring regressors and 6 motion parameters were included as covariates. An F-contrast for the main effect of the N-back task and a t-contrast for the effect of working memory (2-back > 0-back) were defined. Finally, to discover the regions that show effect of working memory, we performed four separate group-level analysis for pre-atDCS, post-atDCS, pre-sham and post-sham sessions. Age and sex were input as covariates.

Eight VOIs were defined as spheres with 6mm radius (Figure 2). The centres of the spheres were in bilateral Inferior Parietal Gyrus (IPG, MNI: −36 −52 43 and 36 −52 43), bilateral Inferior Frontal Gyrus pars triangularis/dorsolateral Prefrontal Cortex (dlPFC, MNI −40 22 27 and 41 31 28), bilateral Superior Frontal Gyrus (SFG, MNI: −22 −1 52 and 28 −2 58), bilateral globus pallidus (MNI: −15 1 4 and 15 5 5). The first eigenvariate of the timeseries of all voxels in each VOI was extracted and adjusted by F-contrast for main effects (removing effects of motion and global signal artefacts). These signals were further used for DCM modelling and analysis.

**Figure 2.**
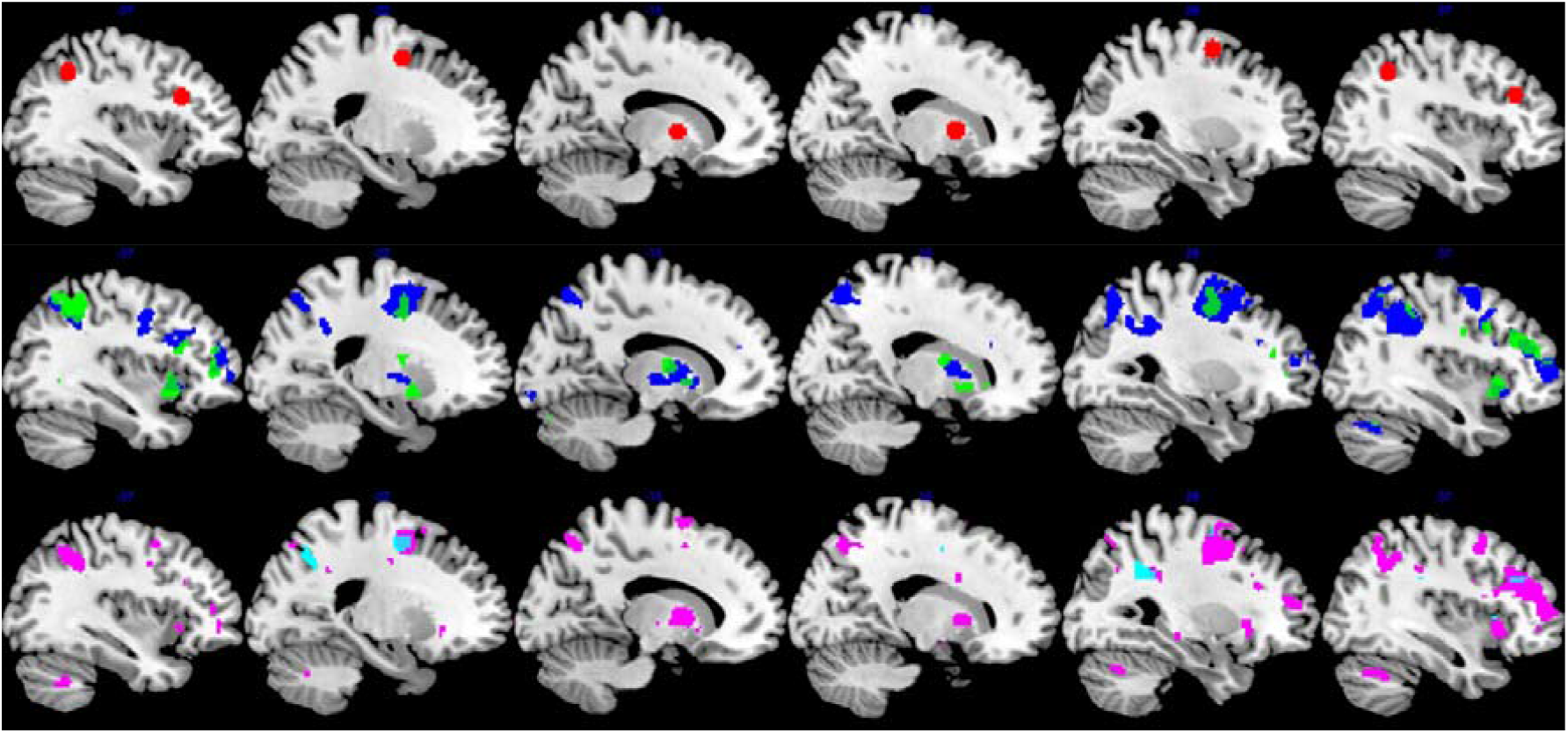
The red spheres signify the location of 8 VOIs selected for the analysis. Clusters signify the uncorrected p <0.001 results of mass univariate 2-back>0-back contrasts performed separately for pre-atDCS (blue), post-atDCS (green), pre-sham (magenta) and post-sham (cyan) sessions.

### Dynamic causal modelling (DCM)

DCM is a generative modelling approach which describes modulating effects of experimental manipulations on effective connectivity [13]. DCMs include average and modulatory connectivity, respectively represented by A-matrix and B-matrix. Average connectivity refers to the average connections between brain regions, independent of task conditions. Modulatory connectivity refers to the impact of experimental inputs on connectivity between regions. The off-diagonal elements of the A-matrix represent the average extrinsic (inter-regional) connection strengths, and the diagonal elements represent the intrinsic (intra-region) inhibitory connection strengths. Meanwhile, the off-diagonal and diagonal elements of the B-matrix represent the modulatory strengths of the experimental manipulation on the average extrinsic and intrinsic connections.

#### First-level DCM

We estimated the effective connectivity for each subject with fMRI DCM [13]. Separate DCMs were constructed and inverted for each session (pre-atDCS, post-atDCS, pre-sham, post-sham). Average connectivity was obtained across all N-back trials in each session. Modulatory effect of working memory was modelled by coding 2-back and 0-back with 1 and −1 respectively. The structure of the full DCMs used is shown in Figure 3. Driving inputs entered the IPG of the two hemispheres [21].

**Figure 3.**
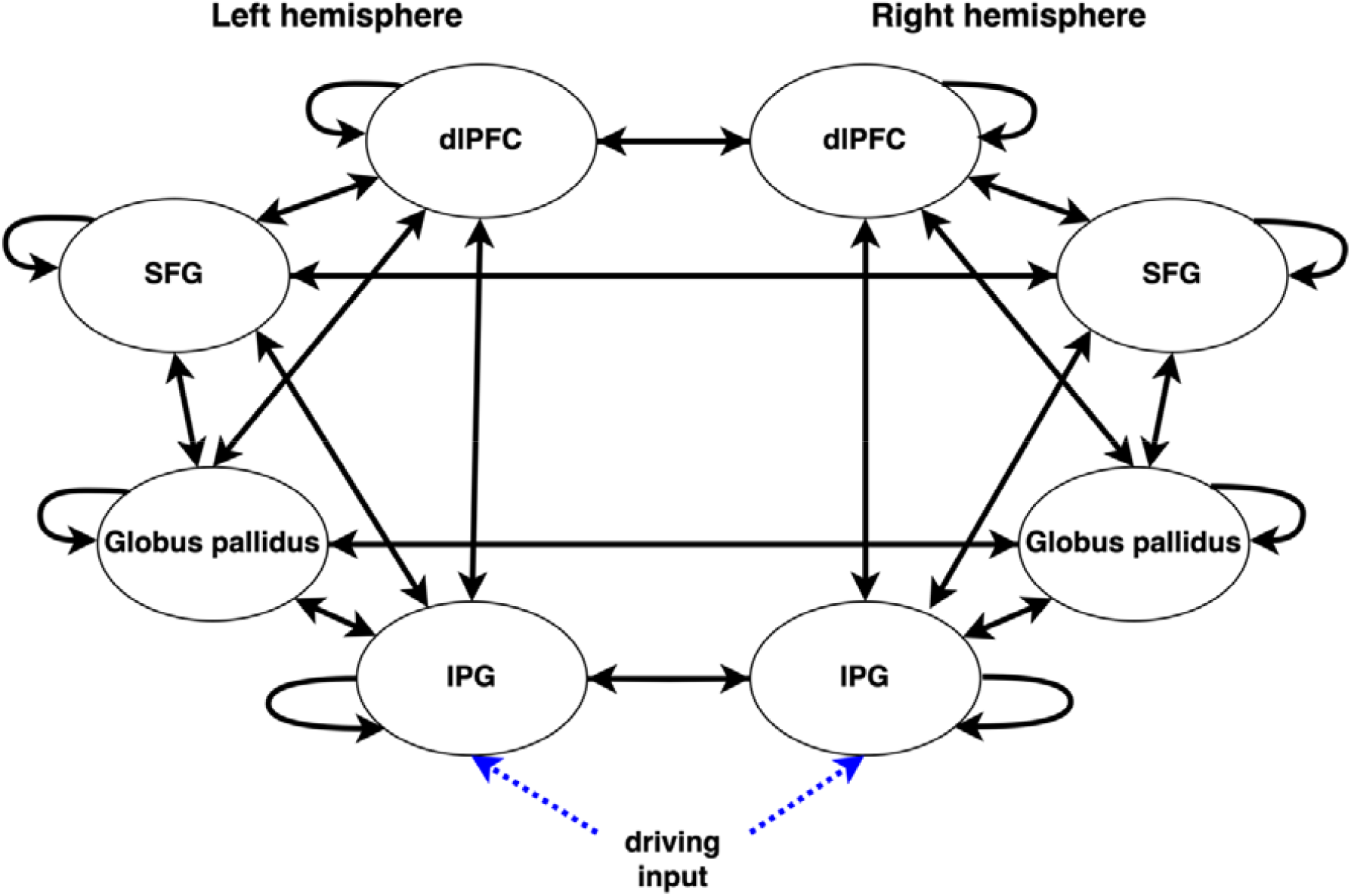
Graph representing the structure of the full DCMs used.

#### Second-level parametric empirical Bayes

A second level-analysis was performed using the parametric empirical Bayes (PEB) approach to identify group level changes in connectivity as result of atDCS stimulation [22, 23]. This model included the mean connectivity, main effect of session, main effect of stimulation type and interaction between session and stimulation type, as well as estimating the between-subject variability. A Bayesian model comparison (BMC) approach was then used to test the hypothesis that interaction between session and stimulation type would manifest in intrinsic and extrinsic connectivity from dlPFC. To do this a model space was constructed by sequentially “switching off” the intrinsic and extrinsic connections originating from dlPFC, producing a total of 16 DCMs. Model comparison between all DCMs in the model space was then carried out based on their model evidences, which were calculated using Bayesian Model Reduction (BMR)[23]. For model comparison, the model evidence was interpreted according to the scales proposed by Kass and Raftery [24].

Finaly, Bayesian model averaging (BMA) was performed to obtain posterior parameter estimates while accounting for model uncertainty. In brief, BMA is an average of the DCM parameter values estimated under each model in the model space, weighted by the posterior probability of each model.

### Relationship between effective connectivity and GABA+ and behavioural performance

For details on the effect of atDCS on GABA+ in the dlPFC and OCC and on behavioural performance, please refer to Garg, Williams [10]. In the present work, Pearson’s correlation analyses were performed to relate how DCM parameters relate to GABA+ from dlPFC only, since no changes related to atDCS were found for OCC.

Next, Pearson’s correlation analyses were performed to relate how DCM parameters relate to behavioural performance. During correlation analysis between average connectivity parameters and behaviour, the average RT and IES for the 0-back and 2-back conditions were obtained per participant, per session. Accuracy was not correlated to average connectivity due to ceiling effects during 0-back condition. During correlation analysis between modulatory connectivity parameters and behaviour, the 2-back accuracy, RT and IES were analysed. In all these analyses, we used a liberal 0.05 alpha for the significance threshold.

## Results

### Session and stimulation effects on effective connectivity

The PEB and BMC methods were used to infer group-level effects of session and stimulation on the average connectivity of 0-back and 2-back conditions. The winning model according to BMC had a posterior probability (Ppost) of 0.33. There was very strong evidence of a positive effect of session on excitatory connections from the left dlPFC to the ipsilateral IPG (Ppost = 0.96), SFG (Ppost = 1) and right dlPFC (Ppost = 1). There was no notable evidence of an effect of session on the connection from dlPFC to the left globus pallidus (Ppost = 0.19). However, there was positive evidence of a positive effect of the interaction between session and stimulation on the excitatory connectivity from left dlPFC to left globus pallidus (Ppost = 0.82), and a negative effect on the excitatory connectivity from dlPFC to the left SFG (Ppost = 0.71). A schematic illustration of these results is presented in Figure 4.

**Figure 4.**
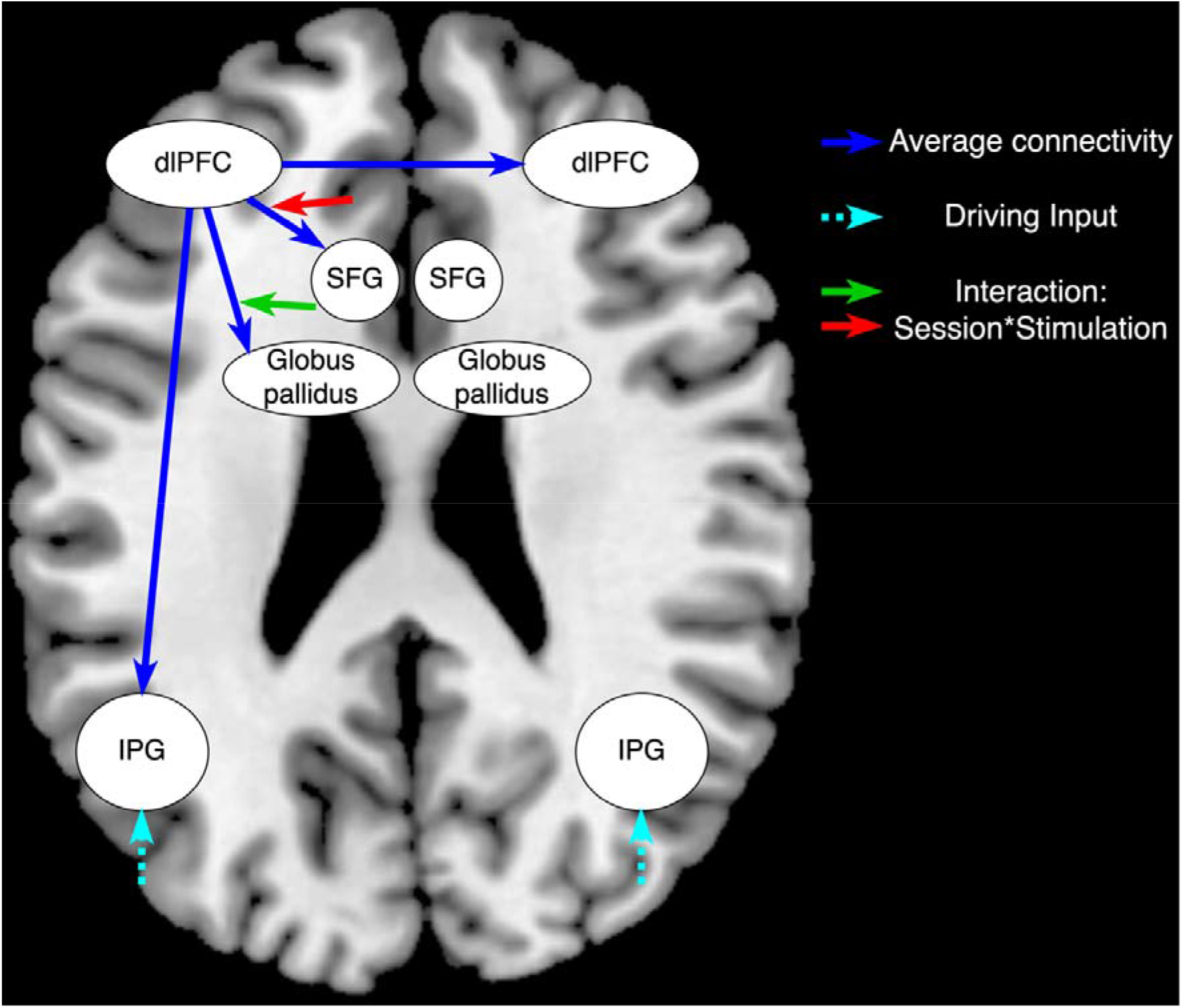
The results of BMA of the BMC winning model. The blue arrows represent the extrinsic excitatory average connectivity from dlPFC; and the green and red arrows represent respectively the positive and negative effect of the interaction between session and stimulation on the extrinsic connections.

PEB and BMC were also used to infer the group level effects of session and stimulation on the modulation strength of working memory load. The null model in which all modulatory strength parameters were turned off was favoured by Bayesian model comparison (Ppost = 0.4), which indicated no effect of session or stimulation on connectivity modulation by working memory load.

### Associations between effective connectivity and GABA+

Following the results of BMC, correlation analyses focused on investigating whether there is any association between GABA+ and left dlPFC’s intrinsic connectivity or its extrinsic connectivity to globus pallidus and SFG (Table 2). There was a significant negative correlation between GABA+ and average intrinsic connectivity of dlPFC during the post-atDCS session (R = −.65, p = .009), such that lower inhibitory intrinsic connectivity was associated with less GABA+. No other associations were found between dlPFC’s connectivity parameters and GABA+.

**Table 2.** The results of correlation analysis between GABA+ and connectivity parameters. Bold font was used to highlight significant results (*p <* 0.05), dashes were placed where no analysis was performed, since there was correlation in either pre-atDCS or post-atDCS session.

|  | pre-atDCS |  | post-atDCS |  | pre- to post-atDCS change |  |
| --- | --- | --- | --- | --- | --- | --- |
|  | R | *p* | R | *p* | R | *p* |
| **Average connectivity** |  |  |  |  |  |  |
| dlPFC to self | -0.25 | 0.331 | **-0.65** | **0.009** | 0.27 | 0.335 |
| left dlPFC to left globus pallidus | 0.14 | 0.594 | 0.26 | 0.349 | - | - |
| left dlPFC to left SFG | 0.1 | 0.695 | 0.18 | 0.515 | - | - |
| **Modulatory connectivity** |  |  |  |  |  |  |
| dlPFC to self | 0.05 | 0.839 | -0.02 | 0.932 | - | - |

### Associations between effective connectivity and behavioural performance

Correlation analyses of average and modulatory pre- and post-atDCS connectivity parameters and behavioural performance revealed no linear correlations between behavioural performance and intrinsic connectivity of left dlPFC or connectivity from left dlPFC to globus pallidus or SFG. Table 3 summarises significant correlations between connectivity parameters and behaviour, and Supplementary Material 4 summarises all performed correlations between connectivity parameters and behaviour.

**Table 3.**
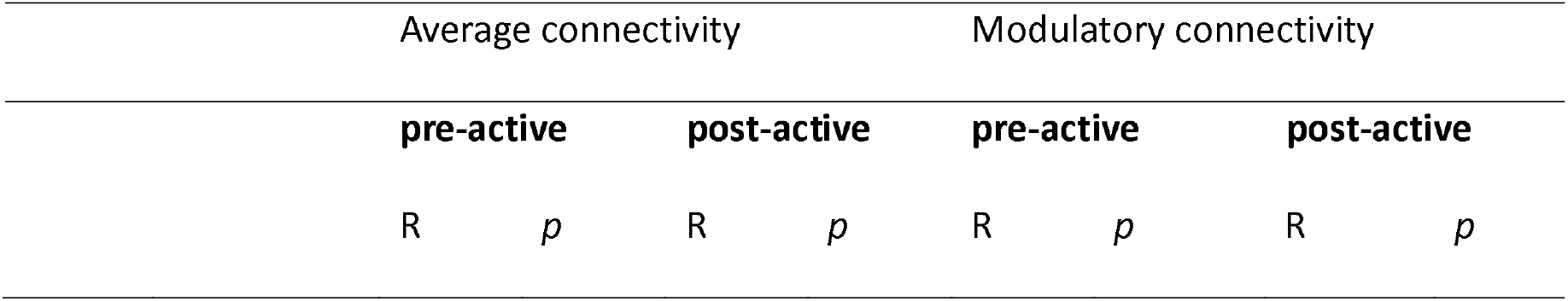
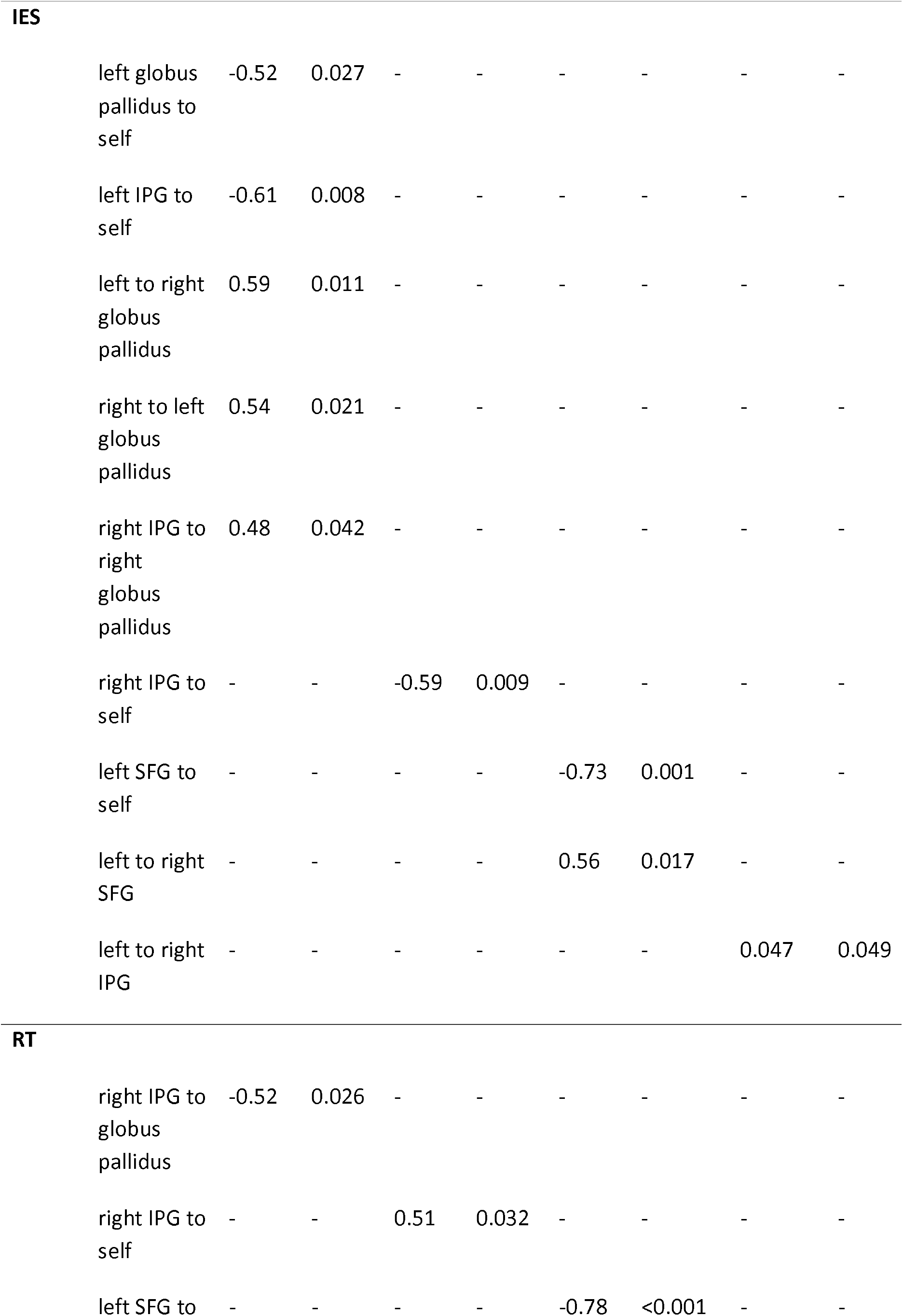
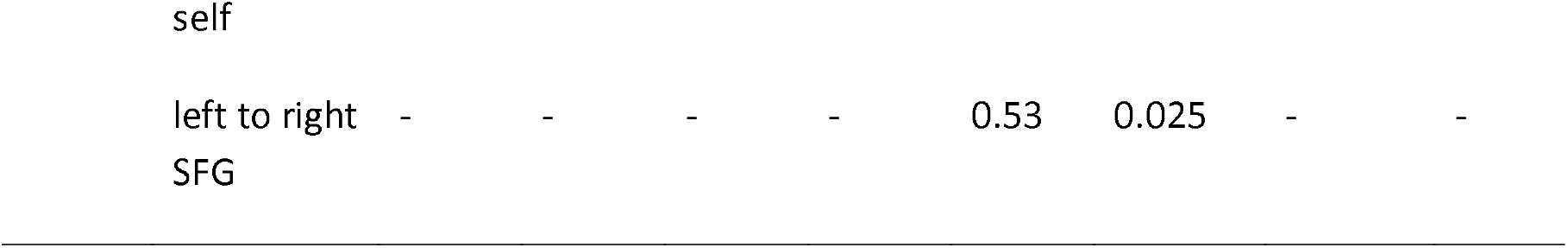
The significant results (*p <* 0.05) of correlation analysis between behavioural performance and connectivity parameters. Dashes were placed where results were not significant.

## Discussion

In this paper, we analysed effective connectivity to investigate the effect of atDCS stimulation on neural correlates of working memory processes. The key finding of this work was that administration of atDCS to dlPFC resulted with reduced connectivity between itself and frontal lobes, coupled with increased connectivity to subcortical structures. This suggests a shift in the functional dynamics of the brain regions involved in working memory and attention. We also discovered neurochemical associations with these changes in the form of an association between GABA+ and sensitivity of left dlPFC to inputs, where reduced self-connectivity of this region was associated with lower GABA+. However, connectivity parameters of dlPFC showed no association with performance on an N-back task. This work offers novel insight to complex interactions between neurostimulation, neural connectivity, and neurochemical processes in the brain. These findings contribute to a better understanding of the role of the left dlPFC in cognitive control functions and open new avenues for research in NIBS and its clinical applications.

This work supplements Garg, Williams [10] with novel analysis of effective connectivity changes associated with administration of sham and anodal atDCS. Here, we uncovered the effects of administration of atDCS that persisted during the post-atDCS session. Specifically, we found reduced the average connectivity from left dlPFC to left SFG and increased connectivity from left dlPFC to left globus pallidus. This finding likely reflects changes to the neural substrates of cognitive processing conducted during N-back task performance. The left frontal regions are implicated in executive control processes through inhibitory control of responses [25], anticipatory attention [26], maintenance of working memory [27], and goal-directed behaviour [28]. Importantly, the left frontal regions have widely been related to error monitoring, interference resolution and selection during retrieval [29-31], particularly but not only in the verbal domain [32]. Therefore, atDCS has likely enhanced excitatory signalling of top-down control and interference resolution of competing stimuli from dlPFC to globus pallidus [33]. Overall, this may result with increased stimulus filtering of relevant stimuli prior to relay of stimuli to working memory processes conducted within the parietal regions [34], or prior to relay of appropriate motor response selection to the sensorimotor cortices [35]. This is an important finding for the NF1 population, because many NF1 patients suffer from increased distractibility and decreased filtering of irrelevant information [36], and as many as 50% of individuals with NF1 receive comorbid diagnosis of Attention Deficit Hyperactivity Disorder that is characterised by difficulty in information filtering [37, 38]. Therefore, the findings suggest that atDCS could potentially serve as a therapeutic tool to address inattention-related symptoms by enhancing the brain’s ability to filter information and improve attentional control. Overall, understanding the effects of these effective connectivity changes on control processes offers important future directions for NIBS research in NF1. Specifically, investigating the specific aspects of cognitive control (e.g., stimulus monitoring, stimulus filtering, sustained attention, selection of motor responses) and specific parameters of atDCS (e.g., intensity, duration, frequency) that yield optimal cognitive benefits for patients could refine treatment protocols.

Further, Garg, Williams [10] demonstrated that application of atDCS was associated with reduction of GABA+. Here we additionally found that lower inhibitory intrinsic connectivity of left dlPFC was associated with less GABA+ post-atDCS, which indicates an increased dlPFC’s excitability during the N-back task post-atDCS. This result suggests that by reducing GABA+, atDCS might make dlPFC more sensitive and responsive to external stimulation. This can affect how the dlPFC processes information, potentially enhancing its ability to engage in working memory performance, attention control, and cognitive flexibility the N-back task. Future research should explore this by investigating how atDCS impacts how regulation of intrinsic connectivity during tasks is related to GABA+ [39-41], and its ratio to glutamate [39, 42].

This study faces several limitations that should be acknowledged. First, DCM BMR did not allow us to test whether the interaction between session and stimulation would be seen in the average intrinsic connectivity of left dlPFC. Instead, DCM BMR always includes all intrinsic connections in the average connectivity. Therefore, we only formally explored the nested models of the effects of atDCS on extrinsic connectivity from left dlPFC. Further, our study is limited by a lack of extensive analysis of psychometrics of attentional capacities in this sample. Therefore, our proposed association between changes to top-down attentional control processes and changes in connectivity and behaviour from pre-atDCS to post-atDCS is speculative. Complementary attentional and linguistic mechanisms in NF1 must continue to be explored to understand the effects of atDCS. We cannot comment on the reliability of long-term effects of atDCS stimulation, and the effects we observed here may be transient. It will also be useful to understand how stimulation of basal ganglia, achieved with other electrode configurations, may impact subcortical signalling during working memory [43].

In summary, this study revealed that atDCS decreased average excitatory connectivity from the left dlPFC to the left SFG and increased connectivity to the left globus pallidus in adolescents with NF1. We suggest that these changes likely reflect enhanced top-down control and filtering of relevant information. Future research should further explore the relationship between attentional processes and neural connectivity in NF1, as well as the long-term effects of atDCS. This study provides novel insights into the neural mechanisms underlying atDCS effects and highlights potential avenues for improving cognitive interventions in NF1 patients.

## Data/code availability

The patient data have been deposited on the Sage Bionetworks data repository https://www.synapse.org/. Approved researchers can request to obtain the data which are subject to data sharing agreements. Codes for data processing and analysis are available at https://github.com/MCLit/NF1-DCM-WM.

## Ethics statement

Ethics approval for the study was obtained from the North West-Greater Manchester South Research Ethics Committee (reference: 18/NW/0762). Written informed consent was obtained from the parents and older adolescent participants and assent was obtained from the younger participants.

## Acknowledgements

This research was supported by the NIHR Manchester Biomedical Research Centre (NIHR203308). ML was funded by the Office for Life Sciences and the National Institute for Health and Care Research (NIHR) Mental Health Translational Research Collaboration, hosted by the NIHR Oxford Health Biomedical Research Centre (NIHR203308). The views expressed are those of the author(s) and not necessarily those of the NIHR or the Department of Health and Social Care. The authors also wish to thank the patients and families that participated in this study. This work was also supported by the Neurofibromatosis Therapeutic Acceleration Program (NTAP) through a Francis Collins Scholarship to SG. JG is supported by NIHR Senior Investigator Award..

