## Supplementary Materials 1-3 for "Effects of non-invasive brain stimulation on effective connectivity during working memory task in Neurofibromatosis Type 1 patients"

#### Supplementary Material 1: Working memory task

The N-back task was used to assess working memory performance in the participants (Kirchner, 1958). Within the scanner, participants were presented with a sequence of black coloured letters on a white screen. The participants were instructed to respond only to the target by pressing a handheld button. During the 0-back condition, the participants responded when the letter ‘X’ was presented on the screen. During the 2-back condition, the participants responded when letter on the screen matched the letter 2 screens before. The stimuli were presented for 2.5 seconds. Each session consisted of 6 blocks of 0-back condition and 6 blocks of 2-back condition, each block was 30 seconds long and consisted of 9 target stimuli. Accuracy was calculated separately for 0-back and 2-back conditions (correct hits + correct omissions/total responses). Response times (RT) were calculated only for time to correct response to target stimuli. Inverse efficiency score (IES) was calculated by dividing RT by accuracy as a measure of speed accuracy trade-off, in which lower scores are generally associated with better cognitive performance (faster response at less accuracy cost) (Bruyer & Brysbaert, 2011).

### Supplementary Material 2: MRS data analysis

Quantification was conducted using the Advanced Magnetic Resonance (AMARES) (Vanhamme et al., 1997) routine in the Java-based magnetic resonance user’s interface (jMRUI5.1, EU project) (Stefan et al., 2009). Individual transients were frequency-aligned, phase-corrected, and averaged within each acquisition session. Water unsuppressed spectra were acquired from the same locations and served as reference. No time-domain filtering was performed on the data before analysis by AMARES. We rejected MRS data from any subject in which there was a change in NAA line width of greater than 3 SD of the global mean linewidth before and after atDCS or sham stimulations such an effect could indicate movement between the two acquisitions. Metabolite concentrations including GABA+, glutamate + glutamine (Glx) and N-acetylaspartate (NAA) were calculated using the unsuppressed water signal from the same voxel as a concentration reference (only GABA+ reported here), without performing a correction for voxel tissue composition. Tissue correction was not considered necessary since the same voxels are interrogated before and after stimulation or sham. In order to perform correlation analysis of GABA+ across subjects, which could be affected by tissue composition differences, MRS voxels were segmented from the T1-weighted anatomical images into grey matter (GM), white matter (WM) and cerebrospinal fluid (CSF) using SPM8 (http://www.fil.ion.ucl.ac.uk/spm/). Voxel registration was performed using custom-made scripts developed in MATLAB by Dr. Nia Goulden, which can be accessed at http://biu.bangor.ac.uk/projects.php.en. The scripts generated a mask for voxel location by combining location information from the Philips SPAR file with orientation and location information contained within the T1 image.

Spectra quality were assessed and artefactual spectra were removed from further analysis, as reported in Garg et al. (2022) and summarised in Table 1. The calculation of partial volume within the VOIs provided the percentage of each tissue type within each voxel (Table 1). There were no significant differences between the percentage of tissue fraction pre-and post-stimulation in any of the voxels. GABA+ was corrected for tissues fraction (GABA/(grey matter + white matter)) for the correlation analyses with effective connectivity.

| Table 1. A summary of removal of spectra from the analysis. Spectra were rejected due to spectroscopic artefacts (e.g. poor water suppression, lipid contamination or broad line widths), > 3SD difference in pre-post intervention NAA line width, and acquisition difficulties. Tissue fraction refers to grey matter + white matter tissues. | | | | | |
| --- | --- | --- | --- | --- | --- |
|  |  | Reason for removal | | |  |
| Session | Spectra location | Spectroscopic artefacts | NAA line width | Acquisition difficulties | Remaining number of spectra |
| Pre-atDCS | dlPFC | 1 | - | - | 28 |
|  | OCC | - | - | - | 29 |
| Post-atDCS | dlPFC | 3 | 2 | - | 24 |
|  | OCC | - | - | 1 | 28 |
| Pre-sham | dlPFC | - | - | - | 31 |
|  | OCC | - | - | 2 | 29 |
| Post-sham | dlPFC | 5 | 1 |  | 25 |
|  | OCC | 2 | 1 | 3 | 25 |

#### Supplementary Material 3: fMRI Processing

Within the present work, we only analyse the fMRI data acquired during pre-stimulation and post-stimulation sessions. Image processing was done using SPM12 (Wellcome Department of Imaging Neuroscience, London; http://www.fil.ion.ucl.ac.uk/spm) and MATLAB R2023a. Dual echo images were extracted and averaged using in-house MATLAB code developed by Halai et al. (2014) (DEToolbox). First, functional images were slice time corrected and realigned to first image. Then, the short and long echo times were averaged for each timepoint. The orientation and location of origin point of every anatomical T1 image was checked and corrected where needed. Mean functional EPI image was co-registered to the structural (T1) image. Motion parameters estimated during co-registration of short echo-time images were input to Artifact Detection Tools (ART; [https://www.nitrc.org/projects/artifact_detect/](about:blank)) toolbox along with combined dual echo scans for identification of outlier and motion corrupted images across the complete scan. The outlier detection threshold was set to changes in global signal 3 z-scores away from mean global brain activation. Motion threshold for identifying scans to be censored was set to 3 mm. Outlier images and images corrupted by motion were censored during the analysis by using the outlier volume regressors. Participants with less than 80% of scans remaining were removed from analysis. Following the removal of participants with high motion and acquisition artefacts, 18 NF1 patients remained. Unified segmentation was conducted to identify grey matter, white matter, and cerebrospinal fluid. Normalisation to MNI space was done with diffeomorphic anatomical registration using exponentiated lie-algebra (DARTEL) (Ashburner, 2007) registration method for fMRI. Normalised images were interpolated to isotropic 2 × 2 × 2 mm voxel resolution. A 6x6x6mm full width at half maximum (FWHM) Gaussian smoothing kernel was applied.
